# Bax mitochondrial residency is more critical than Bax oligomerization for apoptosis

**DOI:** 10.1101/618868

**Authors:** Tomomi Kuwana, Louise King, Katia Cosentino, Julian Suess, Ana Garcia-Saez, Andrew P Gilmore, Donald D Newmeyer

## Abstract

The Bax protein plays an important effector role in apoptosis by forming pores in the mitochondrial outer membrane. While doing so, Bax forms higher-order oligomers in the membrane, but it remains unclear whether this oligomer formation is essential for pore formation. Using cell-free and cellular experimental systems, we investigated two Bax C-terminus mutants, T182I and G179P. Neither mutant formed large oligomers when activated in liposomes. Nevertheless, the G179P mutant could produce membrane pores, suggesting that large oligomers are not required for permeabilization. Surprisingly, however, when G179P was transduced into Bax/Bak double knockout mouse embryonic fibroblasts, it was purely cytoplasmic and failed to mediate cell death. T182I behaved in the opposite manner. When mixed with liposomes, T182I was inefficient in both membrane insertion and permeabilization. However, transduced into cells, BaxT182I resided constitutively in mitochondria, owing to its slow retrotranslocation and mediated apoptosis as efficiently as wild-type Bax. We conclude that Bax’s mitochondrial residence (regulated by targeting and retrotranslocation) is more important for apoptosis than its efficiency of membrane insertion and higher-order oligomerization.

## Introduction

Mitochondrial outer membrane permeabilization (MOMP) is a critical step in the apoptosis pathway. MOMP involves the formation of large protein-conducting pores in the outer membrane. As a result, multiple proteins that are normally restricted to the mitochondrial intermembrane space, such as cytochrome C and SMAC/DIABLO, are released into the cytoplasm, where they trigger the activation of proteases in the caspase family, leading to cell death (Danial & Korsmeyer, 2004). This process is regulated by Bcl-2 family proteins, which can be either pro- or anti-apoptotic and share up to 4 Bcl-2 homology (BH) domains (Chipuk et al., 2010, Gillies & Kuwana, 2014, Youle & Strasser, 2008). Pro-apoptotic Bcl-2 family proteins are further subdivided into two groups; effector molecules (Bax and Bak), which permeabilize the membrane, and BH3 domain-only proteins (Bid, Bim, Puma, Bmf, Bad, Bik, Hrk, Noxa etc.), which promote the activation of Bax and Bak.

An intensive body of research has been directed towards unraveling the molecular mechanisms of MOMP, as this step is considered a ‘point of no return’ in most forms of apoptosis and thus may be a good target for therapeutic intervention (Adams & Cory, 2018). In particular, Bax/Bak activation is of considerable interest for the development of drugs able to kill cancer cells. Cancer cells are more susceptible than normal cells to death following inhibition of anti-apoptotic Bcl-2 family proteins, because their mitochondria are ‘primed’ for death (Certo et al., 2006, Deng et al., 2007, Walensky et al., 2004). This mitochondrial priming phenomenon results from upregulation of pro-apoptotic BH3-only family members along with a compensatory upregulation of anti-apoptotic family members. The pro- and anti-apoptotic proteins form inactive heterodimers in the mitochondrial outer membrane, which serve as reservoirs of the pro-apoptotic BH3-only proteins. Under conditions of apoptotic stress, additional BH3-only proteins are upregulated, displacing the activator BH3-only proteins (e.g. Bim) from the heterodimers. These BH3-only proteins then become available to activate Bax and Bax, leading to cell death. In recent years, BH3-mimetic drugs have been developed that directly disrupt the heterodimeric complexes, thereby inducing MOMP and apoptosis (Oltersdorf et al., 2005). One of these drugs, Venetoclax, is specific for heterodimers containing Bcl-2 and has been approved by the FDA to treat a certain form of chronic lymphocytic leukemia (Cang et al., 2015). In addition, compounds directly activating Bax can also promote MOMP and apoptosis (Reyna et al., 2017). MOMP has therefore been validated as a key process promoting cell death.

Previously, we investigated the molecular mechanisms of MOMP, using systems based on liposomes loaded with fluorescent dextrans, or vesicles comprised of purified mitochondrial outer membranes (outer membrane vesicles, or OMVs), which spontaneously reseal and can also be loaded with fluorescent dextrans (Kushnareva et al., 2012, Kuwana et al., 2002). These vesicles, when mixed with recombinant Bax and a BH3-only protein such as cleaved Bid, can recapitulate the fundamental aspects of the membrane permeabilization process (Kuwana et al., 2005, Kuwana et al., 2002). In further studies using cryo-electron microscopy (cryo-EM), we showed that activated Bax mediates the formation and indefinite enlargement of circular pores in the vesicle membrane, reaching diameters in the hundreds of nanometers (Gillies et al., 2015, Schafer et al., 2009). We later demonstrated that Bax molecules densely line the pore edges (Kuwana et al., 2016). Other groups, using super-resolution microscopy, detected circular arrangements of Bax (presumably corresponding to the pores we saw by cryo-EM) in mitochondria in apoptotic cells (Grosse et al., 2016, Salvador-Gallego et al., 2016).

While Bax exists in large heterogeneous oligomers in these pores (Kuwana et al., 2016), it remains unclear whether large-scale Bax oligomerization is required for pore formation. Indeed, some observations suggest that Bax monomers might be sufficient. First, we showed earlier that the rate of pore formation in native mitochondrial outer membranes is strictly dependent on the concentration of Bax monomers, implying that the rate-limiting step of pore formation involves only the monomeric form of Bax (Kushnareva et al., 2012). Another study used cryo-EM to image nanodiscs containing a single molecule of membrane integrated Bax. The images, produced by averaging multiple nanodiscs, revealed that one Bax monomer can perturb the membrane to form a pore (Xu et al., 2013), arguing for a toroidal (lipidic) pore mechanism.

Now, based on these observations, we have now asked whether Bax oligomerization is necessary for pore formation. Using point mutations in the Bax protein, we experimentally separated different steps in the mechanism of Bax function. Our data show that Bax-induced membrane permeabilization can be uncoupled from higher-order oligomerization. Also, our findings show that efficient Bax-mediated MOMP is critically dependent on the steady-state extent of Bax residency in the mitochondrial outer membrane, prior to activation. The steady-state mitochondrial content of Bax is influenced both by the targeting of Bax to mitochondria as well as the rate of Bax retrotranslocation from mitochondria to the cytoplasm.

## Results

### Membrane insertion of Bax, but not higher-order oligomer formation, is critical for liposome permeabilization

To determine whether Bax oligomerization is required for pore formation, we tested two Bax mutants in the C-terminal transmembrane helix 9 of Bax. The laboratory of Jialing Lin (Zhang et al., 2015), identified one mutant, BaxT182I, as being competent in membrane insertion, but defective in higher-order oligomerization (dimer-dimer interaction). Another study showed that the G179P mutation produced a defect in helix 9 interactions, in studies where the helix 9 sequence was joined to a fusion partner (Andreu-Fernandez et al., 2017).

To test the ability of these mutants to mediate membrane pores, we generated recombinant T182I or G179P mutant Bax proteins and tested them in cell-free vesicle systems. To measure the continuous kinetics of permeabilization, we used an antibody that quenches fluorescence of fluorescein-conjugated dextran molecules once they are released from permeabilized vesicles. At any given time, the fluorimeter thus measures only the signal from dextran molecules residing the vesicles that remain intact. We used the cleaved form of Bid (cBid) to activate Bax species. As shown in Fig. 1A (left panel), Bax 179P was almost as potent as wild-type (WT) Bax in membrane permeabilization. In contrast, BaxT182I protein was much less efficient.

**Figure 1.**
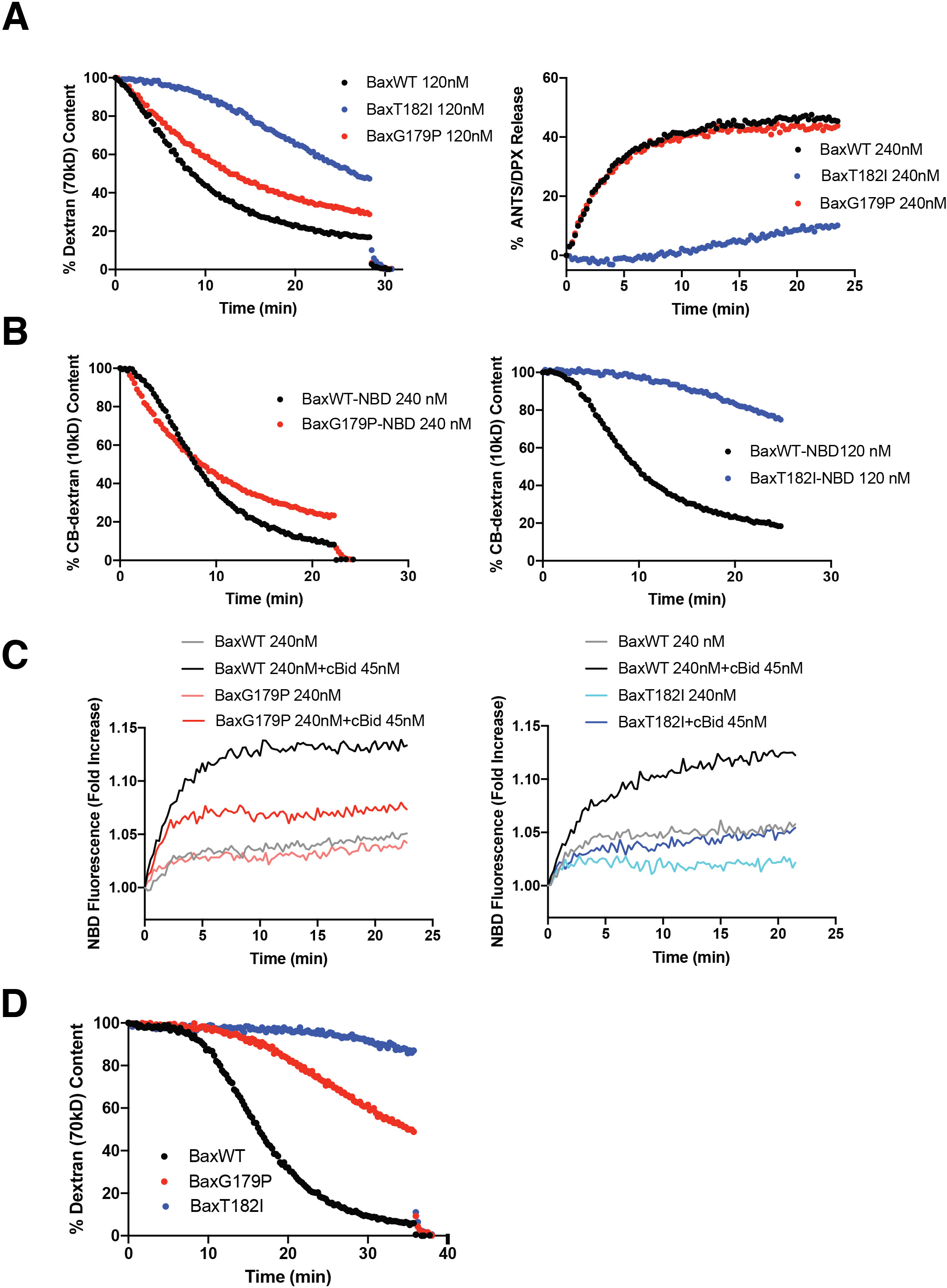
Two Helix 9 mutants of Bax, T182I and G179P, display divergent activities in liposomes. A BaxT182I is less active than WT in liposomes, while BaxG179P is almost as active as WT. Left panel: release kinetics of fluorescein-dextran (70kD) from liposomes were permeabilized with recombinant Bax proteins mixed with the activator, cleaved Bid (cBid)(45 nM). Proteins and liposomes were incubated at 37°C and dextran release was monitored as quenching by anti-fluorescein antibodies. Right panel: release kinetics for liposomes loaded with quenching concentrations of ANTS and DPX (both ~0.5 kD). Bax and cBid were incubated with the liposomes and permeabilization was measured as dequenching of ANTS and DPX resulting from dilution of the dyes once released from the vesicles. B Activities of NBD-labeled Bax species, measured in the presence of cBid (45 nM) on cascade-blue (CB)-dextran (10 kD) loaded liposomes, using anti-CB antibodies. CB-dextrans were used to avoid interference with the spectrally overlapping NBD dye. C BaxT182I is inefficient in membrane insertion. Membrane insertion was monitored using NBD labeled Bax (WT vs G179P in the left panel; WT vs T182I in the right panel) with and without cBid (45 nM) on non-loaded liposomes All the panels (A-C) are representatives of multiple experiments (3 to 5).

An earlier study reported that BaxT182I apparently formed ‘smaller’ pores than WT Bax in isolated mitochondria (Zhang et al., 2015), and we attempted to confirm this in our cell-free system. We loaded liposomes with small reporter molecules, ANTS and DPX, ~500 Da in size. We found that even these small molecules were released only very slowly by BaxT182I (Fig. 1A, right panel). Thus, this result shows that T182I is deficient in the formation of pores of any size.

To determine whether the defect in BaxT182I function occurs at the step of membrane integration, rather than at subsequent events in the pore formation process, we labeled WT and mutant Bax species with nitrobenzoxadiazol (NBD). NBD fluorescence increases if the dye enters a hydrophobic environment, thus signaling that Bax has become inserted into the membrane (Kushnareva et al., 2012, Lovell et al., 2008). First, we verified that NBD labeling did not significantly impair Bax function (Fig. 1B). When we analyzed fluorescence changes upon activation of the Bax variants, we saw that the difference in activity levels correlated with their rate of membrane insertion. Fluorescence of NBD-BaxT182I increased very slowly, compared with WT (Fig 1C, right panel). On the other hand, fluorescence of NBD-BaxG179P increased as rapidly as WT, although it plateaued at a lower intensity (Fig. 1C, left panel). The reason for this is unknown, but we speculate that conformational changes in this mutant may go to completion faster (Kale et al., 2014), or, perhaps that labeling of the cysteine residues in this mutant was inefficient. Regardless, we can say that the difference in membrane permeabilization activity of these Bax species correlates well with the rate of membrane insertion. We conclude that BaxT182I is deficient at the early step of membrane insertion. As liposomes are highly artificial membranes, we investigated the activity of the Bax mutants on OMVs (Kushnareva et al., 2012, Kuwana et al., 2002)(Fig 1D). These native vesicles contain a full complement of mitochondrial outer membrane proteins and thus provide a more physiological membrane environment. The results showed that the response of OMVs to the Bax mutants appeared slower than that in liposomes, but G179P was still more active than T182I.

### Membrane pores can form and enlarge in the absence of higher-order Bax oligomerization

We examined the apparent sizes of mutant and WT Bax oligomers in the membrane using size-exclusion column chromatography (SEC). We incubated liposomes with Bax and cBid, then solubilized them with 1.2% CHAPS detergent (Fig. 2A), a treatment thought to preserve the oligomeric status of Bax (Kuwana et al., 2016). WT Bax formed heterogeneous oligomers that were excluded from Superdex 200 (>600 kD), whereas BaxT182I and BaxG179P oligomers in the membrane, the largest of which reached the excluded volume of Superdex 200 (>600 kD). In contrast, BaxT182I and BaxG179P complexes peaked at an apparent size of 100-200 kD. Bax in solution was predominantly monomeric (~25 kD) with a minor species (possibly dimeric) at around 50 kD, as seen by others (Garner et al., 2016). To measure the precise stoichiometry of Bax oligomers, we used a previously described TIRF microscopy approach (Subburaj et al., 2015). In this method, recombinant fluorescently-tagged Bax (BaxWT-488 or BaxG179P-488) and its activator, cBid, were mixed with unilamellar liposomes. After one hour incubation, these proteoliposomes were fused to obtain a continuous supported lipid bilayer (SLB) containing Bax oligomers (Fig. 2B) on a glass slide. The fluorescence intensities of individual Bax particles (bright spots in the image of Fig. 2B) were then taken to calculate Bax stoichiometry, using monomeric Bax as a calibration reference (Subburaj et al., 2015). We first confirmed that BaxWT-488 and BaxG179P-488 were monomeric in solution (Supplementary Fig. 1). Unfortunately, the labeled T182I mutant aggregated in solution, which precluded the stoichiometry analysis in the membrane. A direct comparison of the intensity distribution of BaxWT-488 (Fig. 2C) with BaxG179P-488 (Fig. 2E) shows lower values of fluorescence intensity for BaxG179P-488, suggesting a lower oligomeric state of particles for this mutant. Importantly, unlike BaxWT-488 (Fig. 2D), BaxG179P-488 was unable to oligomerize into higher-order oligomers at comparable densities in the membrane (Fig. 2F), which is in agreement with the SEC data in Fig. 2A. This conclusion is consistent with reports that Bax helix 9 is involved in higher-order oligomer formation (Andreu-Fernandez et al., 2017, Zhang et al., 2015). Importantly, because BaxG179P could permeabilize liposomes almost as well as WT Bax, we conclude that higher-order oligomers are not required for membrane permeabilization.

**Figure 2.**
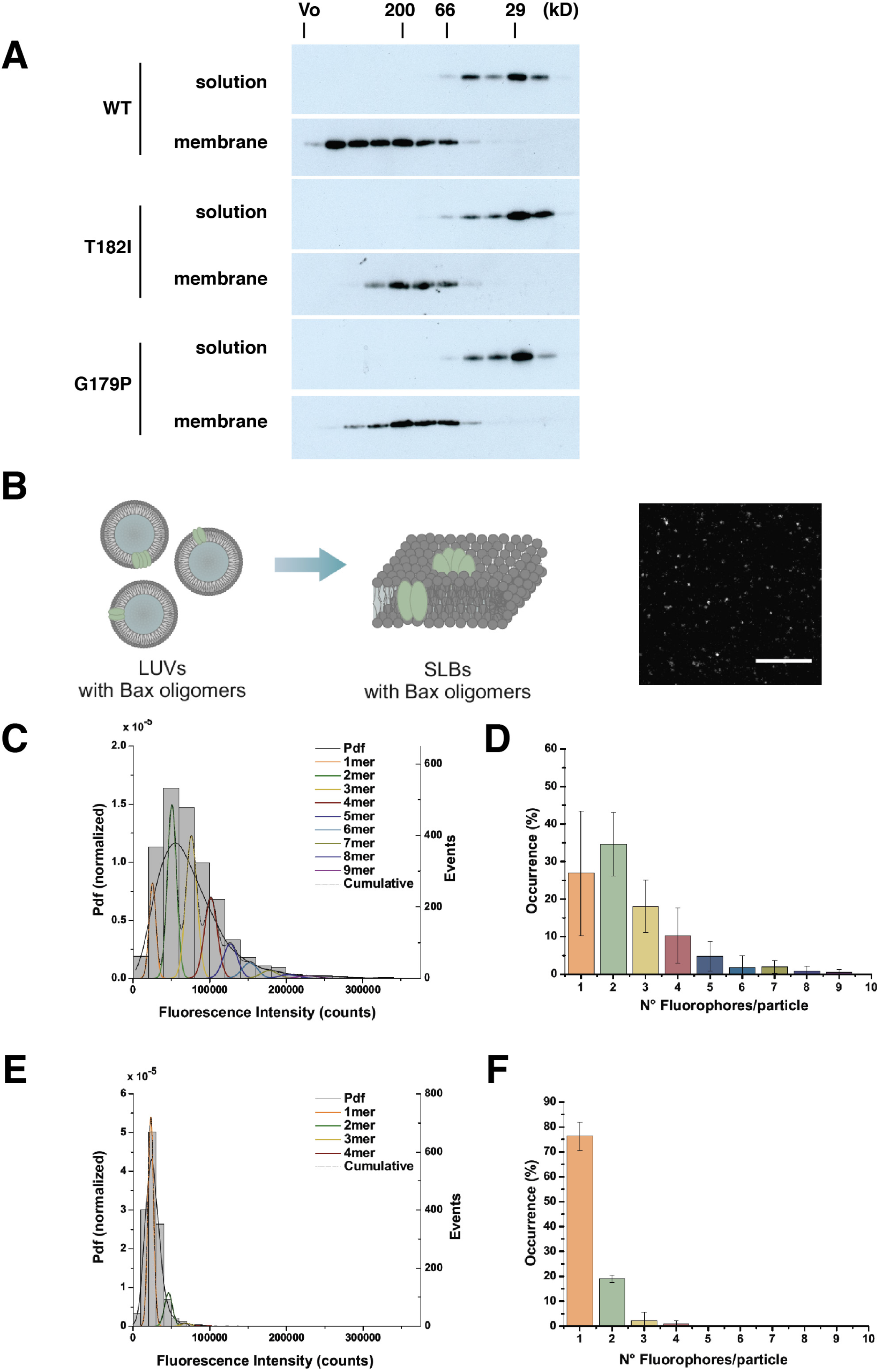
Bax helix 9 mutants form size-restricted oligomers in the membrane. A Size-exclusion chromatography reveals that BaxT182I and BaxG179P from much smaller complexes than WT. Bax species (2 μM each) and cBid (1.3 μM) were incubated with liposomes for 2 h at 37°C. After the incubation period, the membrane fraction was separated in a sucrose gradient centrifugation step to remove unbound Bax protein. Membranes were dissolved in 1.2% CHAPS and fractionated in a Superdex 200 Increase (10/300) column. Fractions were probed with anti-Bax antibody. The data shown are a representative of three independent experiments. B An in situ TIRF–based assay for measuring the stoichiometry of Bax complexes in supported lipid bilayers (SLBs). Schematic representation of the stoichiometry distribution assay on Bax (S4C C62S C126S)-488 and BaxG179P (S4C C62S C126S)-488. LUVs (gray) were incubated with 2.5 nM Bax (light green) and 5 nM cBid (not shown) for 1 h at RT to form proteoliposomes (LUVs containing Bax oligomers). After incubation, these liposomes were used to prepare supported lipid bilayers (SLBs) with labeled Bax oligomers associated with them. The right panel shows a representative TIRF image of a SLB containing Bax oligomers (bright spots). Scale bar is 10 μm. C, E Representative intensity distribution of Bax-488 (C) and Bax G179P-488 (E) mutant particles (minimum 1600 particles per experiment) bound to SLBs prepared from proteoliposomes after 1 h of incubation with the protein. The obtained brightness distribution was plotted as a probability density function (Pdf, black) or, alternatively, as a histogram and fitted with a linear combination of Gaussians to estimate, from the area under each curve, the percentage of occurrence of particles containing n-mer labelled molecules (see color code in the graph). The cumulative fit is shown by a dashed black line. D, F TIRF analysis reveals that BaxG179P fails to form higher-order oligomers in SLBs. Percentage of occurrence of different oligomeric species for Bax-488 (D) and Bax G179P-488 (F) mutant particles calculated as the average value from three (D) and two (F) different experiments. Data provided are the raw values, where no correction for partial labeling (80% for Bax-488 and 50% for Bax G179P-488) was applied (see Materials and Methods). The error bars correspond to the standard deviations from the different experiments.

### Bax helix 9 mutation affects mitochondrial localization

To see the effect of the helix 9 mutations in a more physiological setting, we permeabilized Bax/Bak double knockout (DKO) mouse embryonic fibroblasts (MEFs) with digitonin and incubated them with recombinant Bax and cBid (Fig 3A). Surprisingly, the results were different from what we had observed with liposomes. Although BaxG179P was fairly potent in permeabilizing liposomes, it was unable to promote the release of cytochrome C and SMAC/DIABLO from mitochondria in permeabilized cells. On the other hand, whereas T182I permeabilized liposomes poorly, it was able to release cytochrome C and SMAC/DIABLO from mitochondria in permeabilized cells, although to a lesser extent than observed with WT Bax. We observed that both cytochrome C and SMAC/DIABLO were released to the same extent, and thus our data are inconsistent with those reported by Zhang et al (Zhang et al., 2015), who observed a differential release of cytochrome C and SMAC. Notably, when we recovered the mitochondria-containing pellet fractions from the permeabilized cells, we saw roughly the same (or higher) levels of BaxT182I as WT Bax (bottom strip in Fig 3A), while cytochrome C/SMAC release by BaxT182I was much less than WT. This suggests that higher levels of BaxT182I may be required to produce the same level of permeabilization as with WT. BaxG179P did not measurably associate with mitochondria, which likely explains why this mutant failed to produce cytochrome C/SMAC release.

**Figure 3.**
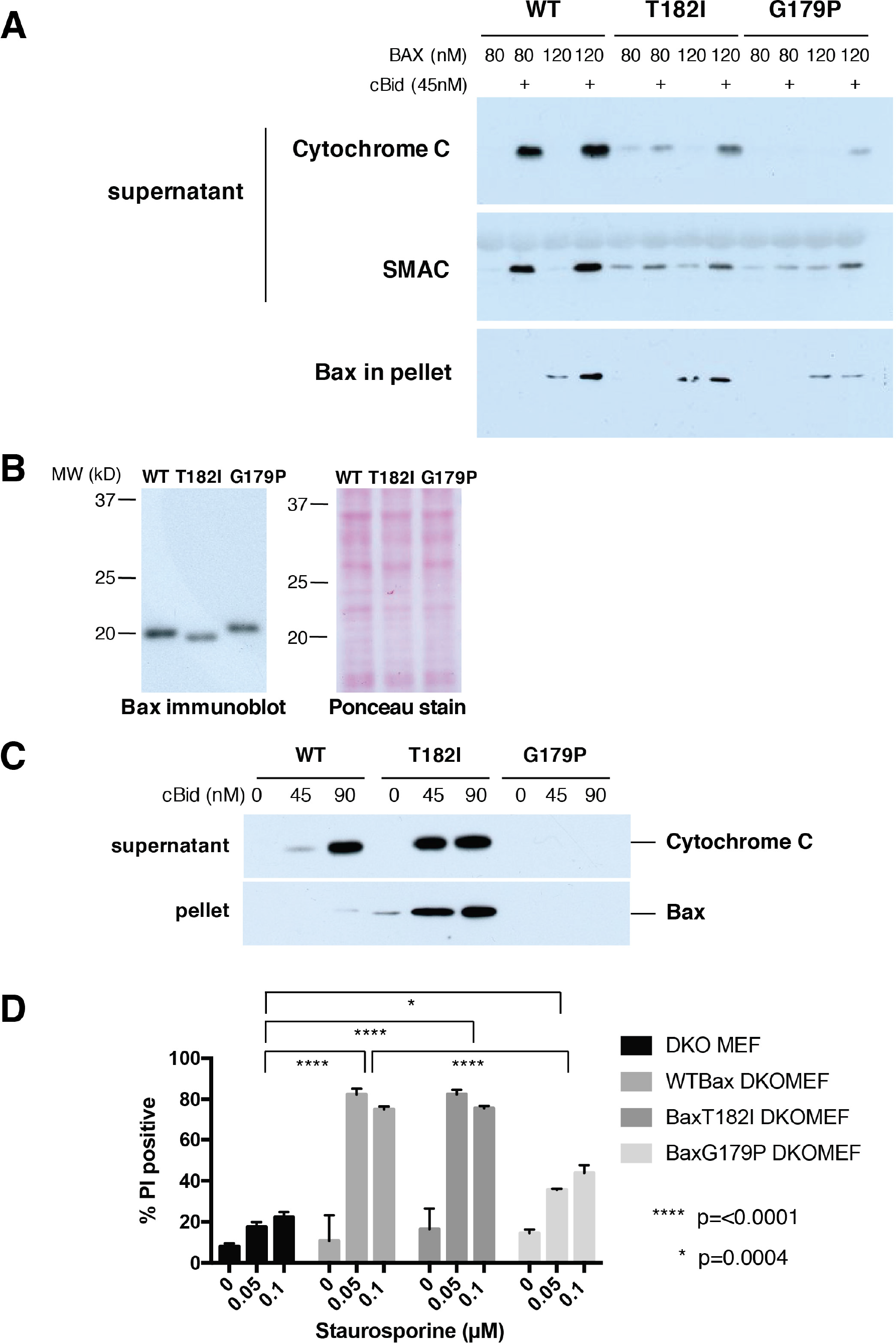
In cells, BaxT182I translocates strongly to mitochondria and, as a result, mediates MOMP and apoptosis as well as wild type Bax; BaxG179P remains cytoplasmic and functions poorly. A Permeabilized cells incubated with recombinant Bax species. Bax/Bak DKO MEFs were permeabilized with digitonin and recombinant Bax proteins and cBid were added for 1 h at 30°C. The supernatant and the pellet fractions were isolated and immunoblotted with anti-cytochrome C, SMAC or Bax antibodies. B Expression of Bax species in transduced cells. Mouse Bax WT, T182I or G179P was transduced in Bax/Bak DKO MEFs using retroviral transduction with IRES-GFP. We sorted out cell populations with equivalent expression levels of GFP, and therefore of Bax species. Note that each mutant Bax migrated slightly different from WT. Ponceau S stain was used to verify equivalent sample loading. C Transduced BaxT182I is recruited strongly to mitochondria in permeabilized cells incubated with cBid. Bax WT, T182I or G179P-transduced DKO MEFs were permeabilized with digitonin and incubated with cBid for 1 h at 30°C. The supernatant and the pellet fractions were immunoblotted with anti-cytichrome C or anti-Bax antibodies. T182I-expressing cells had a higher level of Bax in the pellet than WT and released cytochrome C even more efficiently than WT. Basal BaxT182I levels in the pellet were higher than with WT Bax and levels increased dramatically in the presence of cBid, showing that BaxT182I was recruited to mitochondria more efficiently than WT and cytochrome C was released more than WT. D BaxT182I mediates apoptosis efficiently, but BaxG179P functions poorly. Staurosporine-induced cell death pf transduced cells was measured by propidium iodide (PI) stain. Error bars are SD of duplicate samples. Representatives of 2-4 independent experiments are shown in this Figure.

### Mitochondrial residency of Bax determines cellular sensitivity to apoptosis

To explain the difference between our results with permeabilized cells versus liposomes, we considered that in liposomes, which contain none of the other proteins present in native mitochondrial membranes, Bax must interact with the membrane directly. We hypothesized that cells contain mechanisms to control the mitochondrial localization and retrotranslocation of Bax that are absent in liposomes and that Bax WT and mutants interact differentially with these systems.

To test this hypothesis, we reconstituted Bax/Bak DKO MEFs with WT or mutant Bax, using a retroviral IRES-GFP vector. By fluorescence-activated cell sorting (FACS), we isolated cell populations with equivalent expression levels of GFP, and therefore of Bax species (Fig. 3B). We permeabilized these cells with digitonin, then incubated them in the presence of cBid protein, to activate the endogenously expressed WT or mutant Bax. Finally, we analyzed the resulting release of cytochrome (Fig. 3C). The levels of translocated BaxT182I in the pellet were much higher than those of Bax WT. In contrast, BaxG179P was undetectable in the pellet, suggesting that this mutant failed to associate with mitochondria and was lost during digitonin permeabilization. We conclude that Bax C-terminal mutations can strongly increase or decrease the mitochondrial content of endogenous Bax protein.

Although recombinant BaxT182I was largely inefficient in releasing dextrans and small molecules from liposomes (Fig. 1) and less efficient than WT in releasing cytochrome C from mitochondria in permeabilized cells (Fig. 3A), the mutant protein within transduced cells was as effective as Bax WT in releasing cytochrome C. This presumably results from the increased levels of endogenous BaxT182I in the mitochondria of the transduced cells, offsetting the strongly decreased efficiency of this mutant Bax in membrane insertion.

Next, we tested the ability of WT and mutant Bax species to mediate apoptosis in intact DKO MEFs. Whether these cells were reconstituted with Bax WT or with T182I, they underwent apoptosis to the same extent when treated with staurosporine. On the other hand, cells reconstituted with BaxG179P were as apoptosis-resistant as DKO MEFs (Fig. 3D). To summarize, the results with DKO MEFs were consistent, whether intact or permeabilized. In contrast, the more artificial systems (liposomes and OMVs), produced results that diverged from those seen in cells.

To explain the differences between whole cells and vesicle systems, we hypothesized that whole cells contain systems that control the cytoplasm-mitochondrial distribution of Bax. We fused WT and mutant Bax with GFP and transiently expressed these proteins in DKO MEFs. GFP-BaxT182I was localized preferentially in the mitochondria, as compared with WT Bax, whereas G179P was mostly cytoplasmic (Fig. 4A). These observations were consistent with our immunoblot results using digitonin-permeabilized cells (Fig. 3C). Next, we induced apoptosis in cells expressing WT or mutant Bax and stained the cells with a Bax antibody (6A7). With WT and T182I, we saw activated Bax puncta typical of apoptotic cells, whereas BaxG179P-expressing cells were not stained by 6A7, which specifically recognizes an epitope in Bax that is unmasked after activation (Fig. 4B). With BaxT182I, staining with 6A7 produced a dimmer signal (Fig. 4B) compared with WT, despite the increased mitochondrial content of this mutant (Fig. 3C). This suggests that the T182I mutation impairs Bax activation, consistent with the slower kinetics of liposome permeabilization seen with this mutant (Fig. 1C). Also, we noted that apoptotic puncta formed by BaxT182I were dimmer and smaller than those formed by WT Bax (Fig. 4C). This difference may reflect the smaller complexes formed by T182I (Zhang et al., 2015)(Fig. 2).

**Figure 4.**
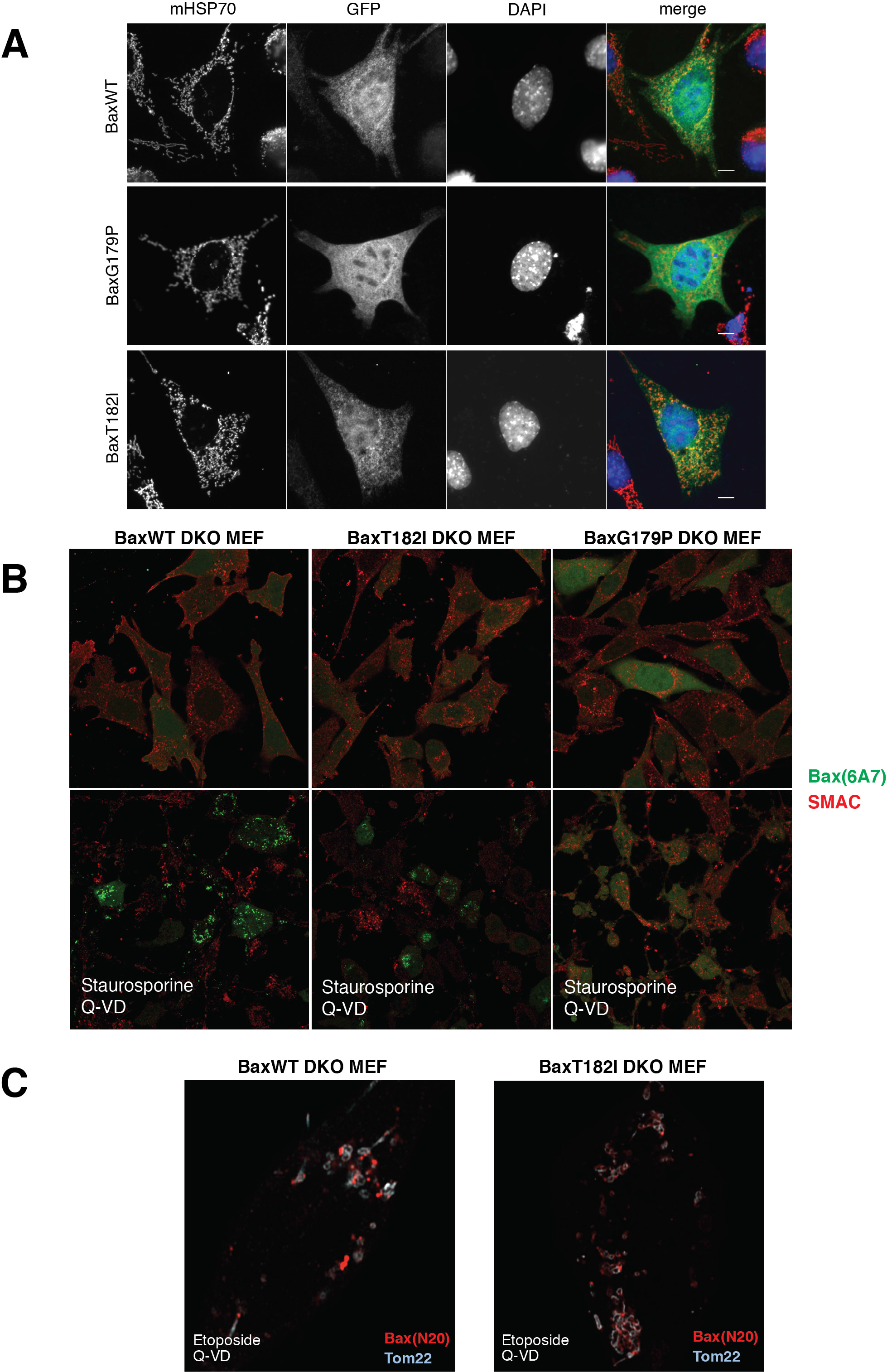
BaxT182I constitutively localizes to mitochondria and forms smaller puncta in apoptosis, whereas G179P is completely cytosolic. A BaxT182I accumulates on mitochondria to a greater extent than WT, while BaxG179P fails to translocate to mitochondria. GFP-Bax was transiently expressed in Bax/Bax DKO MEFs and imaged by epifluorescence microscopy. GFP-Bax was stained with anti-GFP antibody. Mitochondria were stained with mitochondrial HSP70 (mHSP70) and the nuclei, with DAPI. Mitochondria were fragmented as previously reported in these cells (Karbowski et al., 2006), and mitochondrial localization of BaxT182I was slightly more augmented than WT Bax. BaxG179P was diffuse cytoplasmic. B Microscopic detection of activated Bax species in apoptotic cells. Bax/Bak DKO MEFs previously transduced with Bax-IRES-GFP (WT, T182I or G179P) were treated with staurosporine in the presence of caspase inhibitor Q-VD, then fixed and stained with antibodies 6A7 (for the activated form of Bax) and anti-SMAC. The diffuse green GFP fluorescence served as a marker for Bax co-expression. C Bax puncta observed in apoptotic cells. Bax-transduced Bax/Bak DKO MEFs (WT and T182I) were treated with etoposide in the presence of Q-VD. An activated form of Bax was detected by anti-Bax antibody (N20) (red) and the MOM was stained with anti-Tom22 (blue). BaxG179P showed no N20 stain in any of the cells treated with etoposide/Q-VD. Bax puncta in BaxT182I-expressing cells were dimmer and smaller, requiring a higher laser power to detect them.

As BaxG179P was able to permeabilize liposomes, it is apparently functional with regard to the mechanistic steps of membrane integration and pore formation. However, in cells, this mutant is essentially devoid of apoptotic function, as it fails to translocate to mitochondria. We hypothesize that this mutant is unable to interact with the machinery that targets Bax to mitochondria. T182I behaved oppositely, inasmuch as it was inefficient in membrane insertion, but displayed stronger localization to mitochondria. Because T182I mediates cell death as potently as WT, its enhanced mitochondrial residency is apparently enough to compensate for its defect in membrane insertion. Our results argue that there are at least two critical steps prior to pore formation: mitochondrial outer membrane residency (defective in G179P and enhanced in T182I) and membrane insertion (inefficient in T182I). We further suggest that the extent of Bax mitochondrial residency is at least as important for cell death as efficiency in the downstream events such as membrane insertion, symmetric dimer formation, and oligomerization.

### A slow retrotranslocation rate increases Bax levels at mitochondria and sensitizes cells to death

Previous reports have suggested that Bax retrotranslocates, i.e. is in dynamic equilibrium between the cytoplasm and the mitochondria (Edlich et al., 2011, Schellenberg et al., 2013, Todt et al., 2015, Todt et al., 2013). Based on our results, we hypothesized that helix 9 mutations could alter this equilibrium. As BaxG179P was not targeted to mitochondria apparently, we were unable to measure a rate of retrotranslocation for this mutant. However, with WT Bax and BaxT182I, we could assess the rate of dissociation of the corresponding GFP-fusion proteins from mitochondria, using FLIP (fluorescence loss in photobleaching) as described (Schellenberg et al., 2013). In this approach, a cytosolic region of GFP-Bax is first bleached, and the decline of fluorescence in the mitochondria is measured over time as the fluorescent protein re-equilibrates between the mitochondrial and cytoplasmic fractions. For comparison, we included GFP-BaxS184V, which shows reduced mitochondrial retrotranslocation (Nechushtan et al., 1999, Schellenberg et al., 2013, Suzuki et al., 2000). As shown in Fig. 5, both GFP-BaxT182I and S184V were retrotranslocated (dissociated from mitochondria) much more slowly than WT Bax. Analysis of the averaged mobile fraction of each of the Bax variants showed that both T182I and S184V have a significantly larger proportion of protein resident in the mitochondria compared to WT Bax.

**Figure 5.**
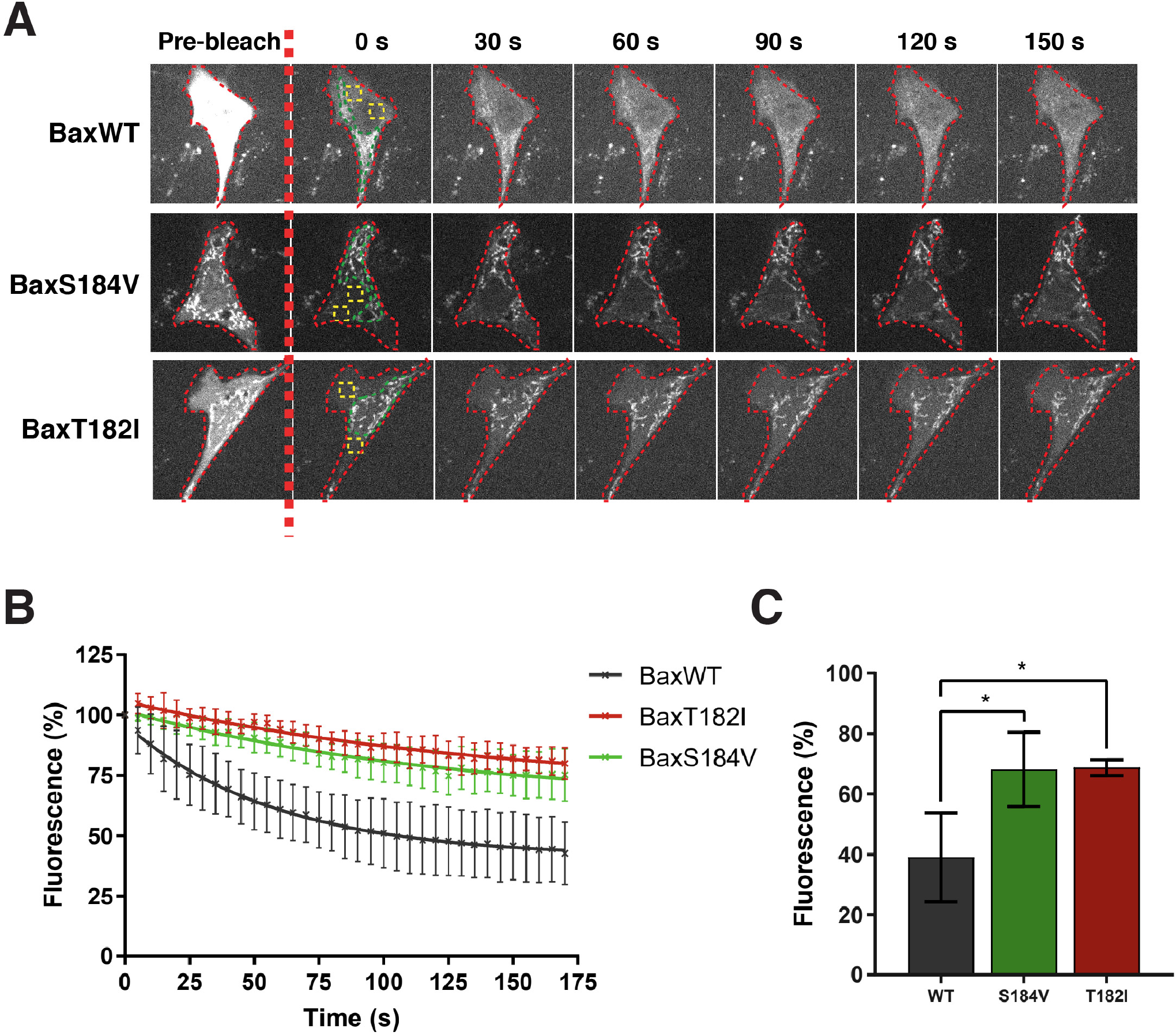
BaxT182I retrotranslocates substantially more slowly than WT Bax. A FLIP analysis of GFP-Bax species transiently expressed in Bax/Bak DKO MEFs. Cytoplasmic fractions of GFP-Bax WT, BaxS184V and BaxT182I were photobleached in the yellow-lined area and fluorescence was monitored in the green ROI. The red line indicates the outline of the imaged cell. B Fluorescence loss of Bax in the measured mitochondrial ROI was normalized to 100% post-bleach and plotted. Data represents average values from 3 independent experiments, 15 cells per experiment. Error bars represent SD. C One phase exponential decay curves were fitted to the data in B and the average mobile fraction calculated from each experiment. Error bars represent SD and data was analyzed via ANOVA. *=p<0.05.

We also developed a permeabilized-cell retrotranslocation assay that is more rapid and convenient than a previously reported method using isolated mitochondria (Lauterwasser et al., 2016). We took digitonin-permeabilized Bax-transduced DKO MEFs and monitored the loss of Bax from the cell pellet over time. We observed that WT Bax was quickly dissociated from the membrane fraction, whereas BaxT182I was much slowly dissociated (Supplementary Fig. 2), consistent with the FLIP assays using live cells.

In summary, our results show that the retrotranslocation equilibrium of BaxT182I is shifted towards mitochondrial residency and that this increased steady-state content of BaxT182I appears to compensate for its defects in downstream events including membrane insertion and higher-order oligomer formation.

## Discussion

We and others have shown that Bax-mediated pores grow to be very large, reaching diameters in the hundreds of nanometers (Gillies et al., 2015, Grosse et al., 2016, Salvador-Gallego et al., 2016). The molecular details concerning how these pores form and enlarge is not well understood. Researchers have commonly assumed that Bax oligomerization is an essential part of the mechanism of pore enlargement. Underlying this assumption is a body of evidence showing that, once Bax and Bak unfold in the membrane, these proteins form symmetric BH3:BH3 groove homodimers that develop into higher-order oligomers through inter-dimer interactions (Bleicken et al., 2014, Dewson & Kluck, 2009, Dewson et al., 2009, Dewson et al., 2008, Dewson et al., 2012). Yet, there has been no direct proof that Bax and Bak oligomers are indeed required for membrane permeabilization.

Symmetric dimer formation of Bax/Bak correlates well with MOMP (Dewson et al., 2008) and Bax is found to exist as even numbered oligomers in supported bilayers, suggesting that dimeric Bax could be the minimum element that is required for permeabilization (Subburaj et al., 2015). On the other hand, our previous work showed that the kinetics of membrane permeabilization are driven by Bax monomers (Kushnareva et al., 2012). Moreover, studies from another group showed by cryo-EM that single Bax molecules inserted into nanodisc membrane could cause a substantial perturbation of the membrane (Xu et al., 2013). These studies suggest a possible poration mechanism involving the concerted action of multiple Bax monomers or dimers, rather than the indefinite growth of Bax oligomers.

Previous reports showed that helix 9 of Bax can mediate inter-dimer interactions that are involved in higher-order oligomer formation (Bleicken et al., 2014, Zhang et al., 2015). To ask whether such oligomers are required for pore formation, we chose to study Bax helix 9 mutations that disrupt inter-dimer interactions while leaving symmetric dimer formation intact, Fortunately, one previous study identified the T182I mutant as fulfilling this criterion (Zhang et al., 2015). We also took note of another study, showing that the G179P mutation abrogated helix 9-helix 9 interactions, in this case when the C-terminal helix 9 sequence was fused to another polypeptide (Andreu-Fernandez et al., 2017).

These two mutations allowed us to interrogate the role of Bax oligomers in MOMP and apoptosis. We found that the disruption of inter-dimer interactions involving helix 9 prevented Bax from forming higher-order oligomers, but surprisingly, did not interfere with membrane permeabilization or cell death. We showed instead that helix 9 plays a significant role in Bax mitochondrial association by affecting mitochondrial targeting (G179P) or retrotranslocation (T182I). We conclude that the steady-state amount of Bax associated with mitochondria can be just as critical in promoting cell death as Bax’s efficiency of membrane insertion and oligomer formation.

### Bax mitochondrial association, but not higher-order oligomer formation, is a critical step in apoptosis

We found that the two helix 9 mutants of Bax (T182I and G179P) exhibited strong phenotypes in cells, corresponding to their mitochondrial localization, rather than their potency in oligomerization or membrane insertion, as measured in model membranes. In particular, we found that BaxG179P is defective in higher-order oligomer formation using two independent approaches: size-exclusion chromatography after CHAPS solubilization (Fig. 2A), and a TIRF microscopic SLB assay (Fig. 2B-F). G179P, despite its defect in oligomerization, could permeabilize liposomes and OMVs and formed pores in liposomes as large as those formed by WT Bax (Fig. 1). BaxT182I displayed a similar inability to form oligomers, as shown by size-exclusion chromatography (Fig. 2A), but unlike G179P, it inserted into the membrane very slowly (Fig. 2C).

In cells, however, these mutants behaved very differently from what we observed with liposomes. G179P was unable to produce MOMP (Fig. 3), apparently because of its defective mitochondrial association (Figs. 3A and 4). We can speculate that the proline residue produces a kink in the helix, dislodging helix 9 somehow aberrantly, which might explain the rapid membrane insertion visible in our liposome assay. However, it appears that this alteration of helix 9 packing entirely abrogates the mitochondrial association of this mutant. This suggests that helix 9 in WT Bax interacts with an unknown Bax targeting machinery in mitochondria and that the G179P mutation disrupts this interaction.

On the other hand, BaxT182I accumulated to greater levels on mitochondria than WT, owing to its slow retrotranslocation rate, and mediated apoptosis as potently as WT Bax. Apparently, the increased concentration of BaxT182I in the mitochondria compensated for its defects in membrane insertion and oligomerization. Another C-terminus mutant, S184V, is known to be located in mitochondria in non-apoptotic cells, presumably due to the disruption of an intramolecular hydrogen bond between S184 and D98; the mutation to valine at 184 is thought to dislodge the helix 9 from the hydrophobic pocket, so that it becomes more available to interact with the mitochondrial outer membrane (Suzuki et al., 2000). In the case of T182I, however, the gain in hydrophobicity from threonine to isoleucine could elicit stronger intramolecular hydrophobic interactions with opposing L76 and/or M79 residues (Dirk Zajonc, La Jolla Institute for Immunology, personal communication). Therefore, it appears that facilitated exposure of helix 9 cannot explain the enhanced mitochondrial localization of BaxT182I. Why this mutant might interact more tightly with the retrotranslocation machinery (i.e. exhibits a slow off-rate) is unclear.

An additional effect of this potentially tighter association of helix 9 with the rest of the molecule is to impede unfolding of the protein, perhaps explaining why we observed a much slower insertion of BaxT182I into the membrane (Fig. 1C). Regardless of the structural mechanisms at play, our data show that the enhanced mitochondrial association of BaxT182I can overcome this mutant’s substantial defect in membrane insertion. We conclude that mitochondrial residency plays an important role in apoptosis induction and oligomer formation.

In this regard, our results are consistent with a previous study from Edlich and colleagues (Edlich et al., 2011). These authors constructed an internally disulfide-tethered mutant of Bax (1-2/L-6), that is constitutively associated with mitochondria due to its slow retrotranslocation rate. Despite the constraints on helix unfolding produced by the cysteine tether, this mutant mediates apoptosis potently, again suggesting that the extended residency of the protein in mitochondrial membranes can compensate for inefficiencies in the downstream steps.

In contrast, another mitochondrially localized mutant, L63E, is unable to mediate MOMP (Kim et al., 2009). Mitochondrial localization of this mutant is suspected to be caused by allosteric helix 9 dissociation, owing to the BH3 helix exposure induced by the mutation. When analyzed by SEC, this mutant elutes at an apparent dimer size, in the presence or absence of tBid, suggesting that the molecule is improperly folded and cannot be activated. In the case of this mutant, it is not surprising that extended membrane residency cannot compensate for an absolute functional defect.

### Bax higher-order oligomers are not required for apoptotic pore formation

Because the Bax mutants we studied here could mediate MOMP without undergoing higher-order oligomerization, we can conclude that large oligomers are not required for apoptotic pore formation. What, then, is the form of Bax that produces pores? One possibility is that Bax monomers are sufficient. An earlier study using cryo-EM demonstrated a single Bax molecule, when activated by a peptide corresponding to the BH3-domain of Bid protein, caused a striking perturbation in the nanodisc membrane into which it became inserted (Xu et al., 2013). Also, in previous studies, we showed that the kinetics of Bax-mediated pore formation are driven entirely by the concentration of Bax monomer (Kushnareva et al., 2012). This implies that monomers are involved in a rate-limiting step in the poration process; however, the data did not rule out the possibility that dimers or oligomers are required in other steps that are not rate-limiting. Finally, another report (Uren et al., 2017) showed that higher-order Bax assemblies can be stabilized by chemical crosslinking, but in the absence of crosslinking they can be disrupted by digitonin treatment. This suggests that higher-order Bax assemblies are dynamic clusters, rather than stable protein chains, admitting the possibility that oligomerization is a byproduct, rather than a cause, of pore formation.

Another possibility is that the permeabilizing unit consists of the symmetric dimer form of Bax, as suggested by previous reports (Subburaj et al., 2015, Westphal et al., 2014). In support of this idea, one recent study (Niu et al., 2017) identified small-molecule inhibitors of Bax-mediated membrane permeabilization that caused the formation of incorrect dimers that could not progress to higher-order oligomers. The authors interpreted the results to mean that monomers are insufficient to form pores. However, because the dimers formed in the presence of inhibitor were not recognized by the 6A7 antibody, it seems possible that the compounds somehow perturb the unfolding process just prior to dimer formation that only aberrant, inactive dimers can be formed.

The hypothetical mechanism we favor is that initially, Bax monomers perforate the bilayer through unfolding of their amphipathic α-helices, in a manner similar to the behavior of antimicrobial peptides in the lipid bilayer (Lee et al., 2008, Lee et al., 2013); subsequently, Bax dimers could play a role in stabilizing the pore walls as they accumulate there, being attracted to their preferred membrane curvature (Uren et al., 2017) and possibly adopting a “clamp” configuration (Bleicken et al., 2014).

### What mechanisms cause Bax mitochondrial localization in cells?

Multiple studies have shown that, in apoptotic cells, Bax and other Bcl-2 family proteins, such as Bak and Bcl-xL, are retrotranslocated continuously from mitochondria outer membranes back to the cytoplasm. The resulting cytoplasm-mitochondria equilibrium regulates the steady-state mitochondrial content of these proteins (Edlich et al., 2011, Schellenberg et al., 2013, Todt et al., 2015). As a result, this equilibrium seems to play a significant role in the cells’ sensitivity to apoptosis. In this study, we have shown that the dynamic exchange of Bax between mitochondria and cytoplasm is influenced strongly by this protein’s helix 9. Based on our data, we postulate that there is a retrotranslocation-related component associated with the mitochondrial outer membrane for which BaxT182I has high affinity (low off-rate) and BaxG179P has low affinity (low on-rate and/or high off-rate).

Parts of the Bax molecule other than helix 9 may also be involved in mitochondrial residency. A recent report described Bax mutations in the so-called non-canonical groove region (helix 1 and 6), where activator BH3-only proteins are thought to interact before engaging the BH3 groove (Dengler et al. 2019). These mutants not only affect the subsequent activation steps, i.e. integration/oligomerization, but also appear to shift Bax equilibrium away from mitochondria at steady state. Therefore, this region is possibly important for interaction with the retrotranslocation machinery, although further studies are required to confirm this.

The molecular mechanisms of retrotranslocation are not completely understood. Both pro- and anti-apoptotic “multidomain” Bcl-2 family proteins (Bcl-xL, Bcl-2, Bax and Bak) participate in it (Edlich et al., 2011), and its rate can be regulated by the activity of protein kinases with known role in cell survival (Schellenberg et al., 2013). One mitochondrial protein important for this process is VDAC2, according to a recent report from the Edlich laboratory (Lauterwasser et al., 2016). VDAC2 has also been shown to interact with inactive forms of Bax and Bak in ways that influence MOMP (Cheng et al., 2003, Lazarou et al., 2010, Ma et al., 2014, Naghdi & Hajnoczky, 2016). The roles of VDAC2 in retrotranslocation and MOMP are therefore legitimate subjects for future study. Because we (Nechushtan et al., 1999, Schellenberg et al., 2013, Valentijn et al., 2003) have observed that the Bax equilibrium significantly influences sensitivity to cell death, it is important to unravel the underlying molecular mechanisms of retrotranslocation, and it is conceivable that the retrotranslocation machinery could prove to be a therapeutic target.

## Materials and Methods

### Recombinant proteins and liposomes

Constructs for BaxT182I and G179P were generated with a QuickChange site-directed mutagenesis kit (#210518; Agilent Technologies, Santa Clara, CA) on the pTYB1 WT-human Bax plasmid (Suzuki et al., 2000) and the sequences were validated. Recombinant non-tagged Bax, BaxT182I, BaxG179P and cBid were generated based on a published protocol (Kuwana et al., 2016, Suzuki et al., 2000). Purity of all the proteins was evaluated to be >90% by Coomassie stained SDS-PAGE. Dextran-loaded or non-loaded large unilamellar vesicles (LUVs) were generated by detergent removal using octylglucoside (OG-LUV)(Schafer et al., 2009). For the Bax oligomerization assay shown in Fig 2, we made larger liposomes by extrusion through a membrane with pore size of 400 nm (Kuwana et al., 2002). Both liposome preparations responded to Bax and cBid the same way.

### Dextran release assays

Bulk kinetic dextran release assays were performed with OG-LUV loaded with fluorescein-dextrans (70 kD)(Sigma; FD70S) or cascade blue (CB)-dextrans (10 kD)(Invitrogen; D1976) and the matched quenching antibodies (Kushnareva et al., 2012). In this assay, the released dextrans from permeabilized vesicles were rapidly quenched by the quenching antibodies present outside the liposomes and the fluorescence decline in real time therefore represented vesicle permeabilization. The percentage of release was calculated against 100% release by 0.2% Triton X-100.

### NBD-labeling of Bax and membrane insertion assay

Recombinant Bax was labeled by NBD dye (IANBD amide, Life Technologies) as reported previously (Kale et al., 2014, Kushnareva et al., 2012, Lovell et al., 2008). 6 μM of Bax and 60 μM of NBD were incubated in 400 μl of labeling buffer (10 mM HEPES pH 7.4, 200 mM NaCl, 0.2 mM EDTA) in the presence of 0.25% CHAPS for 2 h RT in the dark. The sample was eluted from a 10-ml G25 column equilibrated with labeling buffer. The eluted fractions with NBD fluorescence peak (detected by 485 nm^EX^ and 520 nm^EM^) were pooled and concentrated. The typical Bax concentration assayed by BCA was 20-30 μM in ~80 ul. Membrane insertion assays were performed using 720 nM Bax and non-loaded OG-LUVs. Fluorescence at 485^EX^/520^EM^ was measured immediately after NBD-Bax was added to the mixture consisting of 10 mM potassium buffer pH 7.4, 50 mM KCl and 1 mM EDTA together with 45 nM cBid in 100 μl at 37°C in the fluorescence plate reader. The fluorescence increase, which indicates the increased hydrophobicity surrounding the NBD dye, was depicted as a fold increase over the starting value.

### Bax oligomer assay with size-exclusion

The size of Bax oligomers in the membrane was analyzed as previously reported (Kuwana et al., 2016). Bax (2 μM) and cBid (1.3 μM) were incubated with extruded liposomes for 2 h at 37°C. Liposomes were floated up by sucrose step gradient centrifugation to separate them from soluble Bax and then collected in a Microcon filteration device with 0.1 μm pore size. The retained membrane fraction was solubilized in a running buffer consisting of 20 mM HEPES pH 7.4, 150 mM NaCl and 1.2% CHAPS and size fractionated on a Superdex 200 Increase 10/300 GL column (GE Healthcare, Pittsburgh, PA). Fractions were immunoblotted with anti-Bax antibody (Santa Cruz; N20).

### Bax oligomer assay in supported lipid bilayers

#### Bax purification and labeling

Full-length human Bid and single-cysteine, full-length human Bax mutant (S4C, C62S and C126S), as well as Bax mutant (S4C, C62S, C126S, G179P) were expressed in Escherichia coli, purified and labeled with Atto 488 (Attotec, Siegen, Germany) as previously described (Subburaj et al., 2015). Excess of label was removed by size exclusion chromatography, and only monomeric fractions were collected for experiments. The activity of the labelled proteins was measured using a giant unilamellar vesicle membrane permeabilization assay. Labelling efficiency was calculated to be 80% for Bax-488 and 50% for Bax G179P-488 by comparing protein and label concentrations with Bradford, spectrometer and fluorescence correlation spectroscopy (FCS) measurements.

#### Supported lipid bilayers (SLBs)

All lipids were purchased from Avanti Polar Lipids. Lipid mixtures containing egg phosphatidylcholine and cardiolipin in a 80:20 ratio were used. To obtain proteoliposomes, LUVs (100 nm) were prepared as described elsewhere (Flores-Romero et al., 2018, Subburaj et al., 2015). LUVs were incubated with 5 nM cBid and 2.5 nM Bax (S4C C62S C126S)-488 or Bax G179P (S4C C62S C126S)-488 mutants for 1h at room temperature. After the indicated incubation time, labeled-Bax-containing proteoliposomes were diluted to a 1:3 ratio with LUVs in order to be in the single-molecule regime. The resulting solution was immediately used to create SLBs on piranha-cleaned glass slides (0.13–0.16-mm thickness; Menzel) (Subburaj et al., 2015). Unbound proteins and non-fused vesicles were removed by careful washing with buffer and the SLBs were immediately imaged.

#### Microscopy and stoichiometry analysis

All experiments were performed using a modified Zeiss Axiovert 200M epifluorescence microscope using a 488 laser (Ichrome MLE-LFA multi laser, Toptica) equipped with a α Plan-Fluor 100x/1.46 oil objective (Zeiss), a Laser-TIRF 3 Imaging System (Zeiss) and a EM-CCD camera (iXon 897, Andor). Samples were illuminated for 35 ms with a delay time between frames of 25 ms (number of frames 1200) with an intensity of ~ 0.1 kW/cm^2^. Stoichiometry analysis was performed as described elsewhere (Flores-Romero et al., 2018, Subburaj et al., 2015). Data are provided as raw values and no correction for partial labelling was applied, because this maneuver would introduce uncertainty, particularly, with BaxG179P, which had a relatively low labeling efficiency.

### Digitonin-permeabilized cells for cytochrome C and SMAC release and retrotranslocation

Bax/Bak DKO MEFs (~10 million cells) were harvested and permeabilized on ice for 5 min with 200 μg/ml digitonin in digitonin-lysis buffer (DLB; 20 mM HEPES pH7.4, 100 mM KCl, 250 mM sucrose, 5 mM MgCl_2_, 1 mM EDTA, 1 mM EGTA). DLB containing 1% BSA was added to stop the reaction, and the cells were pelleted, washed with DLB and resuspended in DLB. 20 μl of digitonin-permeabilized cell suspension was incubated with recombinant Bax (80-120 nM) with and without cBid (45 nM) at 30°C for 1 h. After the reaction, the cells were pelleted and the supernatant was immunoblotted with anti-cytochrome C antibody (BD; 556433). Then, the membrane was then stripped and probed with anti-SMAC antibody (MBL; JM3298). The supernatant was removed completely from the pellet, which was resuspended in DLB, sonicated briefly and loaded onto SDS-PAGE. Bax was detected by anti-Bax antibody (Santa Cruz; N20). Anti-mouse (Santa Cruz; sc-2005) or rabbit (Invitrogen; 31460) secondary antibodies conjugated to HRP were used to detect primary antibodies.

Bax-transduced MEFs were treated the same way as above and incubated with cBid alone (45 or 90 nM) at 30°C for 1 h. The incubation mix was processed as above for cytochrome C or Bax immunoblotting. For the biochemical retrotranslocation assay, the permeabilized cell suspension was incubated at RT and collected 50 μl at each time point in ice. The supernatant and the pellet of the suspension were processed the same way as above and immunoblotted with anti-Bax, HSP70 (BD; 610607) or VDAC1 (Calbiochem; 31HL) antibodies.

### Transduction of WT Bax, T182I or G179P in Bax/Bak DKO MEFs

Mouse WT Bax, T182I and G179P were cloned into pMIG (a gift from Dr. Emily Cheng) and retrovirus was produced by transfecting Plat-E cells with the plasmid. Bax/Bak DKO MEFs were infected with the virus and the cells were sorted for high level of GFP expression. The cell lysates were prepared in PBS containing 1% NP-40 with protease inhibitors and Bax levels were examined by immunoblotting with anti-Bax antibody (N20). Equal loading was confirmed by Ponceau stain on the transferred nitrocellulose membrane.

### Cell death assay

Bax-transduced MEFs (20,000/well) were seeded into a 96-well plate and treated with staurosporine at 0.05 and 0.1 μM in duplicate for 20 hours. The cells were detached with Accutase (Innovative Cell Technologies, San Diego, CA), washed with PBS in the plate and stained with propidium iodide (Invitrogen)(1 μg/ml) in 100 μl PBS. Propidium iodide fluorescence was detected by FACSCanto II (BD Biosciences, San Jose, CA) in a 96-well plate format.

### Transient expression of GFP-Bax and retrotranslocation assay

Mouse WT Bax, BaxT182 and Bax G179P were cloned in pAcGFP1-C1 (Clontech) and human BaxS184V was cloned in pEGFP-C1 (Clontech). Bax/Bak DKO MEFs were grown in DMEM supplemented with 10% FBS. Cells were seeded onto glass-bottom dishes (IBL) 24 hours before transfection to ensure 60-70% confluency on the day of transfection. Transfections were carried out using Lipofectamine LTX Plus (Life Technologies) following the manufacturer’s instructions, using 0.5 μg DNA. Cells were imaged 24 h post-transfection.

For immunofluorescence, cells were fixed in 4% formaldehyde and permeabilized in PBS/0.1% Triton X-100. Anti-GFP antibodies (Life Technologies; A-11122) were diluted to working concentration in PBS/10% horse serum (37°C for 1 h). After washing in PBS, secondary antibodies were diluted in the above buffer (37°C for 1 h). After washing in PBS, DAPI was diluted to 1 μg/ml in PBS and incubated for 5 min. Coverslips were mounted using Dako Fluorescence Mounting Medium (Agilent), and imaged using an Axioplan 2 imaging microscope with a 63x 1.4NA Plan Apochromat objective (Zeiss MicroImaging) and quantified with Image J software.

Live cell imaging was carried out using a Zeiss Axio-Observer Z1 microscope with CSU-X1 spinning disc confocal (Yokagowa), using a 63X/1.40 Plan-Apochromat objective, Evolve EMCCD camera (Photometrics) and a motorized XYZ stage (ASI). The 488nm laser was controlled via AOTF through the laserstack (Intelligent Imaging Innovations). Images were captured using Slidebook 6.0 software (Intelligent Imaging Innovations). FLIP was carried out by photobleaching two regions within the cell, one cytosolic and one overlapping the nucleus, for two 50 ms iterations, 100% laser power. Images were captured every 5 s. Captured images were analyzed using Image J. In brief, the background was subtracted and the cytosolic content exclusive of the bleached ROIs was selected. Signal decay was quantified and normalized to 100% fluorescence post-bleaching. Statistical analysis was carried out by ANOVA, using GraphPad Prism.

### Confocal microscopy

WT Bax, BaxT182 or BaxG179P-transduced Bax/Bak DKO MEFs were plated out in 8- chamber coverglasses (Lab-Tek II Chambered coverglass #1.5; 155409) and treated with 1 μM staurosporine and 20 μM Q-VD (Selleckchem) for 5 h. Cells were fixed with 0.5% glutaraldehyde and quenched with 0.5% sodium borohydride in PBS. After permeabilization with 0.5% TritonX100, cells were stained with anti-Bax antibody (6A7; Biolegend) and anti-SMAC antibody (Sigma; PRS2411) at 1:100, with 10% BSA in PBS. Primary antibodies were detected by anti-rabbit antibody conjugated to Alexafluor (AF) 568 and anti-mouse antibody to conjugated to AF488 (Life Technologies). The cells were imaged in a Zeiss LSM780 microscope.

For imaging in the Airyscan, WT Bax or BaxT182I-transduced Bax/Bak DKO MEFs were plated out on coverslips and treated with etoposide (400 μM; Selleckchem) and Q-VD (20 μM; Selleckchem) for 16 h. Cells were fixed and permeabilized as above and stained with anti-Bax antibody (Santa Cruz; N20) and anti-Tom22 antibody (Sigma; 4G4) at 1:100, then with the secondary species specific antibodies conjugated with AF568 (for Bax) or AF647 (for Tom22)(Life Technologies). The coverglasses were mounted with ProLong Gold (Life Technologies) and inspected with Zeiss LSM880 with a 63x 1.46 NA alpha Plan-Apochromat objective. The images were processed with Airyscan FAST (Zeiss).

## Acknowledgements

We thank Dr. Emily Cheng (Memorial Slone-Kettering Institute, NY) for providing the pMIG vector and the protocols for retroviral transduction. The Zeiss LSM880 with airyscan was acquired by NIH S10OD021831 to the microscopy core (LJI). This work was supported by the Manchester Cancer Research Centre by a CRUK training award (L.K.), a core grant from the Wellcome Trust (203128/Z/16/Z) to The Wellcome Center for Cell-Matrix Research (A.P.G.), the Eliteprogramme for Postdocs of the Baden-Wurttemberg Stiftung (K.C.), ERC-St12 309966 (A.G-S), DFG GA1641/2-2 (A.G-S) and NIH R01CA179087 (D.D.N.).

## Author contributions

T.K. conceived the project. T.K., L.K., K.C., J.S., A.G-S., A.P.G. and D.D.N. designed and performed experiments and analyzed the data. T.K. and D.D.N. wrote the manuscript with the input from all the authors.

---

**Supplementary Figure 1. Labeled WT and G179P Bax species are monomeric in solution**

A Schematic representation of the protocol used for sample preparation in B and C. LUVs (grey) were used to prepare SLBs. SLBs were incubated with 2.5 nM labeled Bax (light green) and 5 nM cBid (not shown) for 30 min or 1 h at room temperature, washed carefully with buffer and immediately imaged by TIRF microcopy.

B, C Percentage of occurrence of fluorophore units per particle for Bax (S4C C62S C126S)-488 (B) and Bax G179P (S4C C62S C126S)-488 (C) mutants directly added on SLBs calculated as the average value from two different experiments. Due to unspecific interaction of Bax molecules with the SLB glass support, Bax molecules are unable to oligomerize, resulting mainly in monomers and some dimers. Data provided are the raw values, where no correction for partial labeling was applied. The error bars correspond to the standard deviations from two different experiments.

**Supplementary Figure 2. Biochemical retrotranslocation assay shows slow mitochondrial dissociation of BaxT182I**

Bax/Bak DKO MEFs transduced with WT Bax or BaxT182I were digitonin-permeabilized and incubated at RT over time. Supernatant and pellet fractions were isolated at each time point and Bax and other proteins were probed by immunoblot. VDAC1 was shown as loading controls for the pellet. A cytosolic protein, HSP70, was leaked from the permeabilized cells, but the level was unchanged over time.

